# Fidgetin-like 2 is a novel negative regulator of axonal growth and can be targeted to promote functional nerve regeneration after injury

**DOI:** 10.1101/2020.03.19.999508

**Authors:** Lisa Baker, Moses Tar, Guillermo Villegas, Rabab Charafeddine, Adam Kramer, Olga Vafaeva, Parimala Nacharaju, Joel Friedman, Kelvin P. Davies, David J. Sharp

## Abstract

The microtubule (MT) cytoskeleton plays a critical role in axon growth and guidance. Here, we identify the MT severing enzyme fidgetin-like 2 (FL2) as a negative regulator of axonal regeneration and a potential therapeutic target for promoting neural regeneration after injury. Genetic knockout of FL2 in cultured adult dorsal root ganglion (DRG) neurons resulted in longer axons and attenuated growth cone retraction in response to inhibitory molecules. Given the axonal growth-promoting effects of FL2 depletion *in vitro*, we tested whether the enzyme could be targeted to promote regeneration in a rodent model of peripheral nerve injury. In the model used in our experiments, the cavernous nerves (CN) are either crushed or transected, mimicking nerve injury caused by radical prostatectomy (RP). As with patients, CN injury results in erectile dysfunction, for which there are presently poor treatment options. At the time of injury, FL2-siRNA or control-siRNA was applied to the site using nanoparticles or chondroitin sulfate microgels as delivery agents. Treatment significantly enhanced functional nerve recovery, as determined by cavernosometry (measurements of corporal blood pressure in response to electrostimulation of the nerve). Remarkably, following complete bilateral nerve transection, visible and functional nerve regeneration was observed in 7 out of 8 animals treated with FL2-siRNA. In contrast, no control-siRNA treated animals showed regeneration. These observations suggest a novel therapeutic approach to treat peripheral nerve injury, particularly injuries resulting from surgical procedures such as RP, where treatments depleting FL2 could be applied locally at the time of injury.

## Introduction

The MT cytoskeleton is a major regulator of axon growth and guidance. It serves as the major track for vesicular transport through the axon shaft, provides structural support to the axon, and, in the distal region of the axon, polymerizing MTs play a critical role in promoting axon elongation and in steering the growth cone in response to environmental cues [1, 2]. Given its role in regulating axonal growth, targeting the MT cytoskeleton to enhance axon regeneration after nerve injury has been proposed as a potential therapeutic strategy [3–11]. This proposition is supported by studies demonstrating that modulation of MT dynamics in the axon can promote regeneration and attenuate degeneration: for example, multiple studies have found that low doses of MT stabilizing drugs (taxol and epothilones) promote axon regeneration and improve recovery of locomotor function after spinal cord injury (SCI) in rats [5–7, 11]. Increasing the density of dynamic MTs in the axon by knocking down the MT severing enzyme fidgetin was also shown to promote axon regeneration after dorsal root crush in rats [9].

MT severing enzymes, which are members of the <u>A</u>TPases <u>A</u>ssociated with diverse cellular <u>A</u>ctivities (AAA+) superfamily, cause breakages in MTs by forming hexameric rings around the C-terminal tails of tubulin and using energy from ATP hydrolysis to pull on the tails, thereby causing tubulin dimers to dissociate from the MT lattice [12, 13]. Through their severing activity, they regulate MT length, number, and branching, and fine-tune the dynamics of the MT cytoskeleton [14]. MT severing enzymes include katanin, spastin, and the fidgetin family (fidgetin, fidgetin-like 1 (FL1), and fidgetin-like 2 (FL2)). With the exception of FL2, all have been reported to play important roles in regulating axonal growth through their remodeling of the axonal MT array [9, 15–19].

We recently identified FL2 as a negative regulator of cell migration and a potential therapeutic target for promoting wound healing of cutaneous wounds: application of nanoparticle-encapsulated FL2-siRNA to murine cutaneous burn and punch biopsy wounds significantly enhanced the rate of wound closure [20, 21]. *In vitro,* the acceleration of cell motility observed with FL2 knockdown was accompanied by an increase in the density of dynamic MTs, particularly at the leading edge of the cell, where FL2 was shown to strongly localize [20]. It was therefore proposed that the enzyme normally suppresses forward movement of the cell by selectively paring down dynamic MTs near the cell cortex. Beyond these two studies, FL2’s function at both the cellular and physiological level are unknown.

The goals of the present study were to characterize the role of FL2 in regulating axonal regeneration and assess its potential as a therapeutic target for promoting regeneration after nerve injury. For the latter, we utilized a rat model of peripheral nerve damage in which the cavernous nerves (CN) of rats are crushed or transected, a model commonly used to investigate treatments to address nerve injury associated with radical prostatectomy (RP) [22, 23]. The CN are parasympathetic nerves which travel along the posterolateral prostate and innervate the penis, where they regulate erectile function by controlling blood flow to the corporal tissue. Because they lie on the prostate, they are highly susceptible to damage during RP resulting in erectile dysfunction (ED) [24].

This model is highly clinically relevant, since currently there are no clearly effective treatments for CN injury-induced ED post-RP [25]. Prostate cancer is the second most prevalent cancer in men [26], and RP is the most common treatment for localized prostate cancer, particularly for younger men who are sexually active [27]. Unfortunately, even with the advent of nerve-sparing procedures, incidences of ED post RP are high: 60% of patients experienced selfreported ED 18 months after RP, and only 28% of men reported erections firm enough for intercourse at a 5-year follow-up according to a prostate cancer outcomes study [25, 28]. ED post-RP has a major impact on the lives of many patients and their partners.

We demonstrate that FL2 is a novel negative regulator of axon regeneration that appears to suppress axonal growth by selectively severing dynamic MTs in the distal axon shaft and growth cone. We further show that FL2 depletion attenuates the effects of inhibitory environmental cues on growth cone advancement. Finally, we demonstrate that targeted knock-down of FL2 after CN injury is effective in promoting regeneration and restoration of nerve function in rats. These observations suggest a novel therapeutic approach to treat peripheral nerve injury, particularly injuries resulting from surgical procedures such as RP, where treatments depleting FL2 could be applied locally at the time of injury.

## Results

### FL2 suppresses axonal growth in dissociated adult DRG neurons

A FL2 conditional knockout (KO) mouse with a tdTomato reporter gene insertion was generated by Genoway (Fig. 1A, see also Figs. S1–S5 and Methods for more information). To test whether FL2 regulates axon growth, adult DRG neurons harvested from FL2-flox homozygous mice were transduced *ex vivo* with adenovirus (AV) containing either a Cre recombinase plasmid (Cre AV) to excise the FL2 gene, or GFP control plasmid (GFP AV). The neurons were cultured for one week to enable sufficient time for FL2 depletion following Cre-mediated excision of the gene. Transduction efficiency was approximately 80% based on the percent of GFP-positive control neurons 5 days after transduction. Because no antibody against rodent FL2 exists, successful gene excision was confirmed by quantification of FL2 and tdTomato mRNA levels one week after transduction by reverse transcription quantitative polymerase chain reaction (RT-qPCR) (Fig. 1B). One week after transduction, neurons were replated at low density to allow neurite growth to begin anew and fixed 48 hours later for morphometric analysis. FL2 depleted neurons had significantly longer axons (indicated by the length of the longest neurite on each neuron) compared to controls, with a mean axon length approximately 50% longer than GFP-AV treated neurons (p<0.0001, Welch’s t-test) (Fig.1C,D). These results were consistent with preliminary studies we conducted in rat neurons transduced with plasmids encoding scrambled or FL2-shRNA and replated for regrowth (Fig. S6). No other differences in neuron morphology (such as branching or growth cone morphology) were observed.

**Figure 1.**
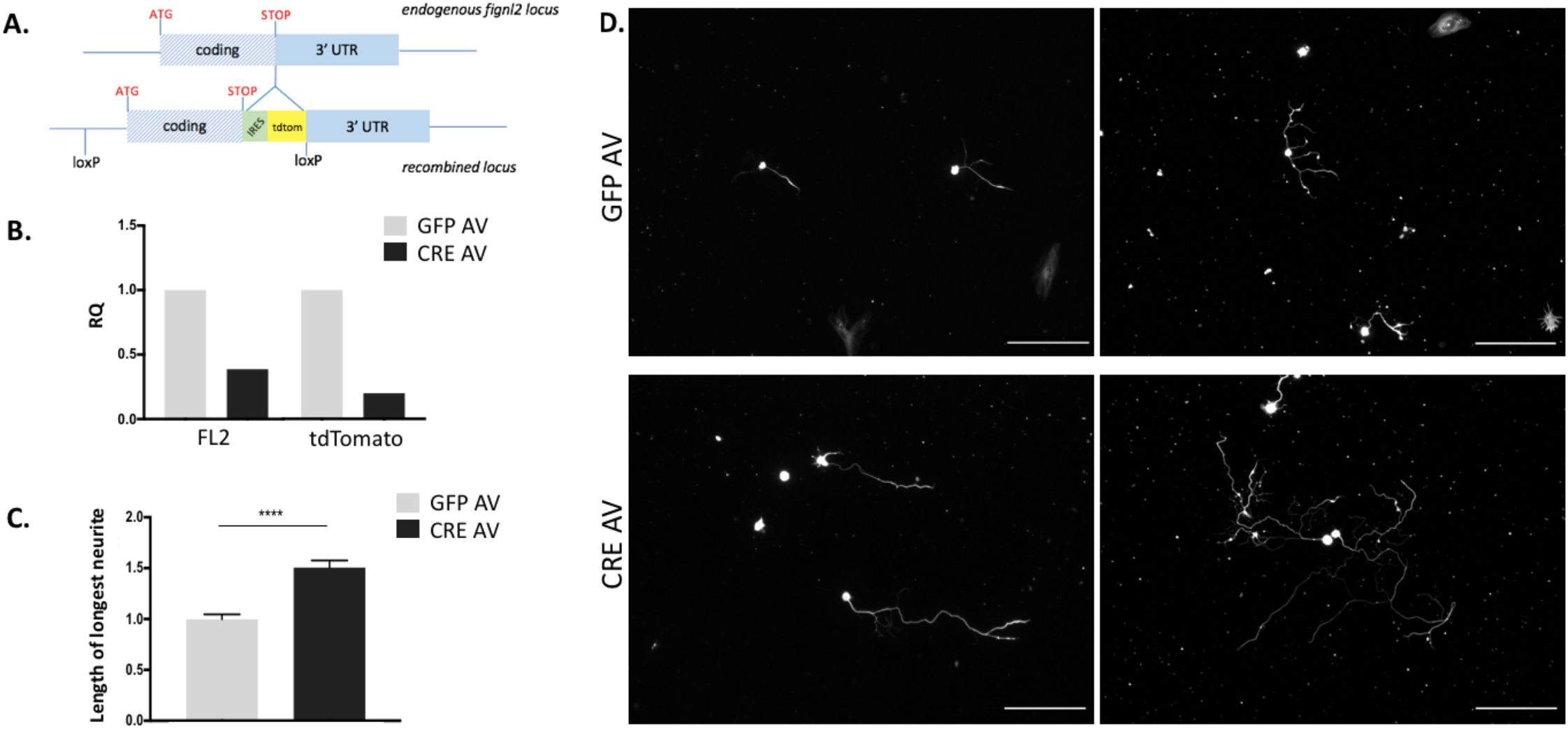
FL2 depletion accelerates the rate of axon regeneration in adult DRG neurons. **A)** Schematic of IRES-tdtomato knock-in and lox sites at FL2 endogenous locus. The FL2 gene is on chromosome 15 and is intronless, composed of 1 exon extending over 4.2 kB. An internal ribosome entry site and tdtomato reporter gene were inserted after the stop codon between the coding sequence and 3’ UTR sequence. LoxP sites were inserted upstream of the start codon and after the reporter gene. **B)** FL2 and tdtomato mRNA levels 1 week after transduction with GFP AV or Cre AV. RNA combined from 4 transduced cultures. **C)** Average length of longest neurites in control and FL2 knockout neurons 2 days after replating (GFP AV: 1 ± 0.05; Cre AV: 1.51 ± 0.07. Mean ± SEM, p<0.0001, Welch’s t-test. GFP AV, n = 517, Cre AV n = 479). **D)** Micrographs of control (upper images) and Cre AV DRG neurons (lower images) replated at low density one week after viral transduction, fixed 2 days later and immunostained for βIII tubulin (scale bar = 200 μm).

### FL2 depletion results in a more dynamic MT array in the distal axon

Axonal MTs are comprised of domains that differ markedly in their stability properties: there is a fraction of labile, highly dynamic MTs, a fraction of stable, long-lived and relatively undynamic MTs, as well as a hyper-stable fraction of polyaminated MTs (referred to as the cold-stable fraction) [29, 30]. Longer-lived (stable) MTs accumulate post translational modifications, such as acetylation, polyglutamylation, and de-tyrosination, while labile MTs are relatively unmodified [31]. It has been hypothesized that an increase in the density of labile, unmodified MTs in the distal axon is conducive to axon regeneration [29, 32, 33], and MT severing enzymes are known to selectively target different MT subpopulations within the axon. For example, katanin and spastin preferentially target stable MTs [34, 35], while fidgetin severs dynamic MTs [19]. We therefore characterized changes in the relative ratios of dynamic and stable microtubules in DRG neurites with FL2 depletion.

To characterize changes in the axonal MT array with FL2 depletion, we measured changes in the ratios of dynamic and stable MTs to total MTs along neurites by quantitative immunofluorescence. Neurons were dual-stained with either tyrosinated tubulin and βIII tubulin or acetylated and βIII tubulin (tyrosinated and acetylated tubulin are commonly used as markers of labile and stable MTs, respectively, while βIII is a neuron-specific isoform of tubulin that is present in both stable and labile MTs). We found that FL2 depleted neurons had significantly higher ratios of tyrosinated to βIII tubulin and a lower ratio of acetylated to βIII tubulin in the distalmost region of neurites: the tyrosinated:βIII tubulin ratio increased by ~26% while the acetylated: βIII tubulin ratio correspondingly decreased by ~26% in the distalmost 50 μm of neurites (p<0.0001, Welch’s t-test) (Fig. 2A-D). The ratios of dynamic and stable microtubules in the mid-proximal neurite shaft were unaffected by FL2 knockout (Fig. 2F), suggesting that the enzyme’s activity and/or localization may be most concentrated in (or restricted to) the distal axon and growth cone.

**Figure 2.**
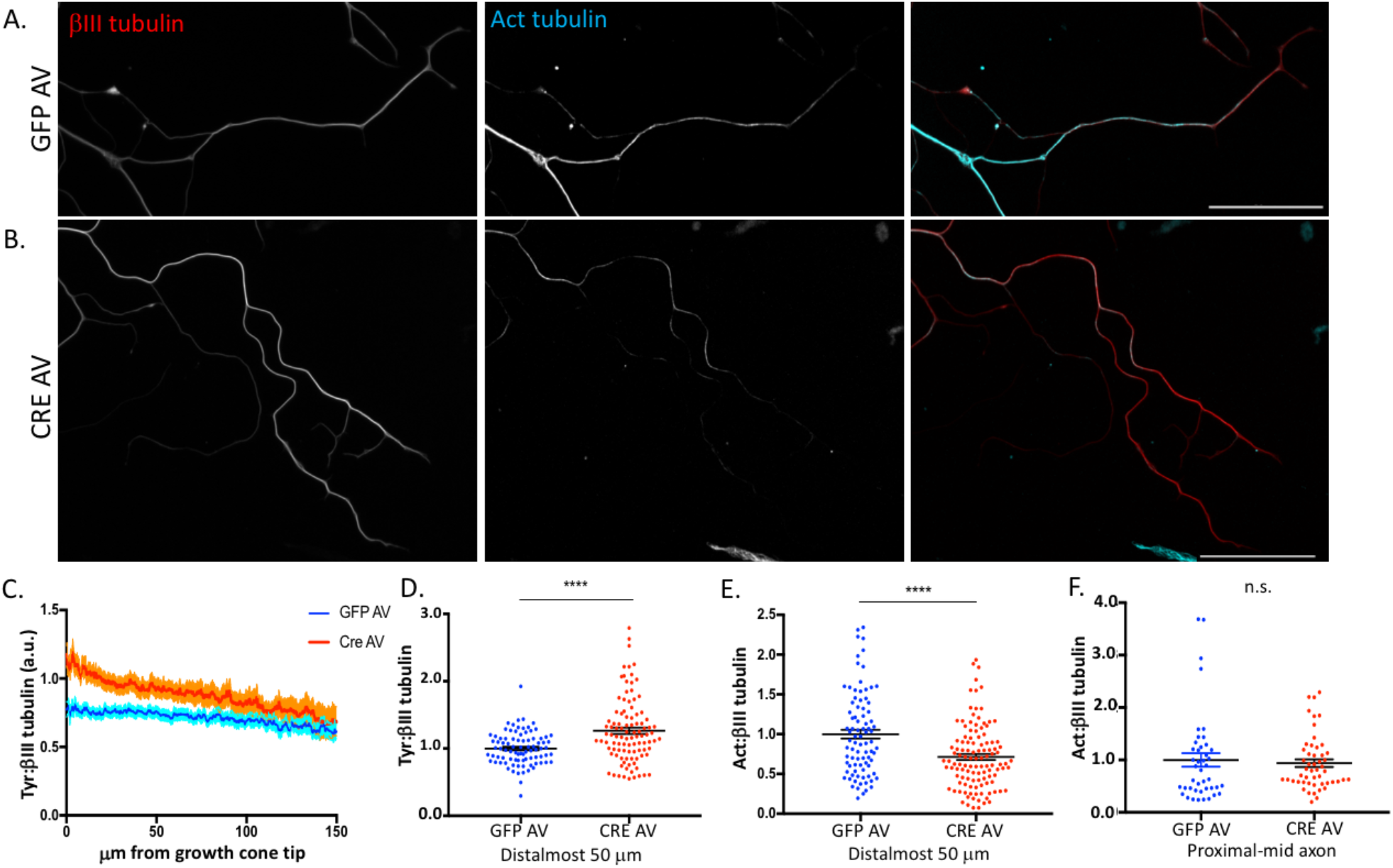
FL2 depletion results in a more dynamic microtubule array near the growth cone in regenerating neurites. **A,B)** Immunomicrographs of adult DRG neurites of GFP AV and Cre AV treated neurons 2 days after replating, which were dual-stained for acetylated (cyan) and βIII tubulin (red). Scale bar = 50 μm. **C)** Traces of the average ratios of tyrosinated to βIII tubulin in GFP and Cre AV treated neurons from one representative experiment, starting at the tip of the growth cone and moving toward the soma. The dark line is the mean intensity, the shaded area represents the SEM for each point along the neurite. **D)** Distribution of the ratios of tyrosinated to βIII tubulin fluorescence intensities in the 50 distalmost μm of neurites, normalized to the GFP AV mean ratio. Each dot represents a single neurite (GFP AV: 1 ± 0.026, Cre AV: 1.26 ± 0.048, p<0.0001, Mean ± SEM, Welch’s t-test, GFP AV n = 86, Cre AV n = 101). **E)** Distribution of the ratios of acetylated to βIII tubulin fluorescence intensities in the 50 distalmost μm of neurites, normalized to the GFP AV mean ratio (GFP AV: 1 ± 0.056, Cre AV: 0.74 ± 0.038. Mean ± SEM p<0.0001, Welch’s t-test, GFP AV, n = 87; Cre AV n = 116). F) Distribution of the ratios of acetylated to βIII tubulin in the proximal to mid region of the axon shaft.

### FL2 mediates growth cone response to inhibitory environmental cues

Given the evidence that FL2 was suppressing axonal growth by severing dynamic MTs in the distalmost axon and growth cone, we hypothesized that FL2 might also mediate growth cone steering. To test this, we challenged regenerating axons with a mixture of chondroitin sulfate proteoglycans (CSPG). CSPG are abundant in the glial scar tissue at central nervous system (CNS) injury sites, and impede axon regeneration through the scar by causing growth cones to retract [36]. We used a modified version of a conventional stripe assay [37] in which neurons were replated onto stripes of CSPG mixed with Alexa-fluor-568-anti-mouse IgG antibody (in order to visualize the stripe boundaries). The majority of adult DRG neurons turn upon encounter with these aggrecan stripes, whereas stripes of Alexa-fluor-568-anti-mouse IgG alone elicit no turning response from growth cones. After 3-4 days, the neurons were fixed and imaged, and the percent of neurites crossing the stripe borders out of total neurite/border encounters was quantified. FL2 depleted neurons had a significantly higher percentage of neurites crossing through stripe borders compared to GFP AV treated control neurons (43%+/-3% versus 26%+/-4%) (Fig. 3A-C).

**Figure 3.**
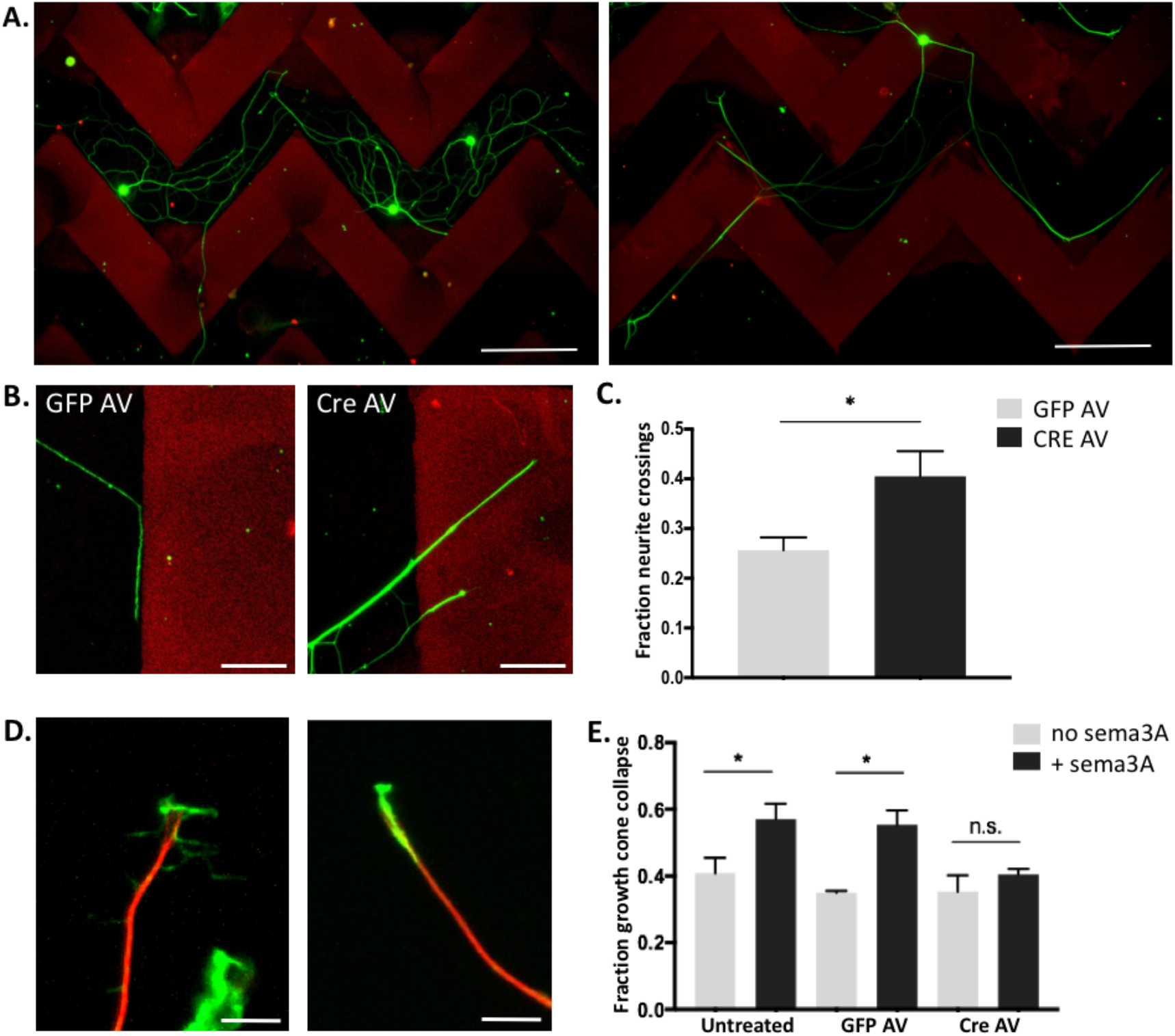
FL2 depletion attenuates the effects of inhibitory substrates on adult DRG growth cone advancement during regeneration. **A)** Lower magnification of GFP AV and Cre AV treated neurons 72 hours after plating on aggrecan stripes. Scale bar = 200 μm. **B)** High magnification image of a GFP AV treated neurite turning in response to an aggrecan border, and Cre AV treated neurites crossing through the stripe border. Scale bar = 50 μm. **C)** Fraction of neurites of GFP and Cre AV treated neurons that crossed aggrecan stripe borders (GFP AV mean = 0.26 ± 0.026; Cre AV mean = 0.41 ± 0.06. p=0.0169, Welch’s t-test, GFP n = 65 neurons, 257 neurite/border encounters; Cre AV n = 79 neurons, 200 neurite/border encounters. Experiment performed 4 times). **D)** Immunomicrographs showing an active, viable growth cone (left) and a collapsed growth cone (right), stained for microtubules in red and actin in green. Scale bar = 10 μm. **E)** Fraction of collapsed growth cones in untreated, GFP AV treated, and Cre AV treated neurons, in the presence and absence of semaphorin3A. GFP and Cre AV experiments performed in quadruplicate, untreated +/− sema3A performed in triplicate (untreated, no sema3A: 0.38 ± 0.04; untreated, +sema3A: 0.56 ± 0.03; GFPAV, no sema3A: 0.35 ± 0.01; GFPAV, +sema3A: 0.55 ± 0.04; Cre AV, no sema3A: 0.35 ± 0.05; Cre AV, +sema3A: 0.40 ± 0.02. Mean ± SEM, p*<0.05, Welch’s t-test).

In a complementary experiment, neurons were challenged with recombinant semaphorin 3A (sema3A). Sema3A is an important chemo-repellent during the development of the nervous system [38, 39], and has also been implicated in axon guidance during adult PNS regeneration [40–42]. *In vitro,* Sema3A causes growth cone collapse in both embryonic and adult DRG neurons [43]. In adult DRG neurons, the Sema3A-induced growth cone collapse phenotype is less pronounced. This is because adult DRG neurons have smaller growth cones that are more likely to assume a bullet-shape—meaning a higher percentage of growth cones exhibit a “collapsed” phenotype to begin with—and, additionally, response to Sema3A is limited in adult DRGs to nociceptive neurons (which comprise the majority of small-bodied DRG neurons), whereas in embryonic DRGs the response to sema3A is ubiquitous [44]. We first tested whether Sema3A induced growth cone collapse in our nine day old cultures of adult DRG neurons two days after replating. We found that treatment with a recombinant Sema3A-Fc chimera caused a ~20% increase in the percent of growth cones exhibiting a collapsed phenotype compared to neurons treated with Fc alone (Fig. 3E), consistent with a previous study which found that growth cones of adult rat DRG neurons with a collapsed phenotype rose 18% with Sema3A treatment *in vitro* [44]. Similarly, GFP-AV treated neurons showed a ~20% increase in growth cone collapse with Sema3A treatment (Fig. 3E). In contrast, FL2 depleted neurons showed no significant increase in growth cone collapse in the presence of Sema3A (Fig. 3E). Together, these results indicate that FL2 plays an important role in mediating growth cone response to inhibitory substrates in adult sensory neurons.

### Targeted knockdown of FL2 after cavernous nerve transection promotes functional nerve regrowth

Based on the growth-promoting effects of FL2 depletion on regenerating axons *in vitro,* we reasoned that using siRNA to knockdown expression of FL2 at the site of nerve injury would promote nerve regeneration and recovery of nerve function. To test this hypothesis, an animal model of nerve injury was used in which the cavernous nerve (CN) is transected, resulting in a gap of 3-4 mm between the proximal and distal nerve segments. At the time and site of injury, animals were treated with FL2-siRNA or control-siRNA embedded into hardened chondroitin sulfate microgels, termed FL2-siRNA-wafer or control-siRNA-wafer, respectively (Fig. S7A). The animals treated with FL2-siRNA-wafer had reduced FL2 expression in the MPG compared to animals treated with control-siRNA-wafer even 14 days after treatment (approximately 43% lower levels) (Fig. 4A).

**Figure 4.**
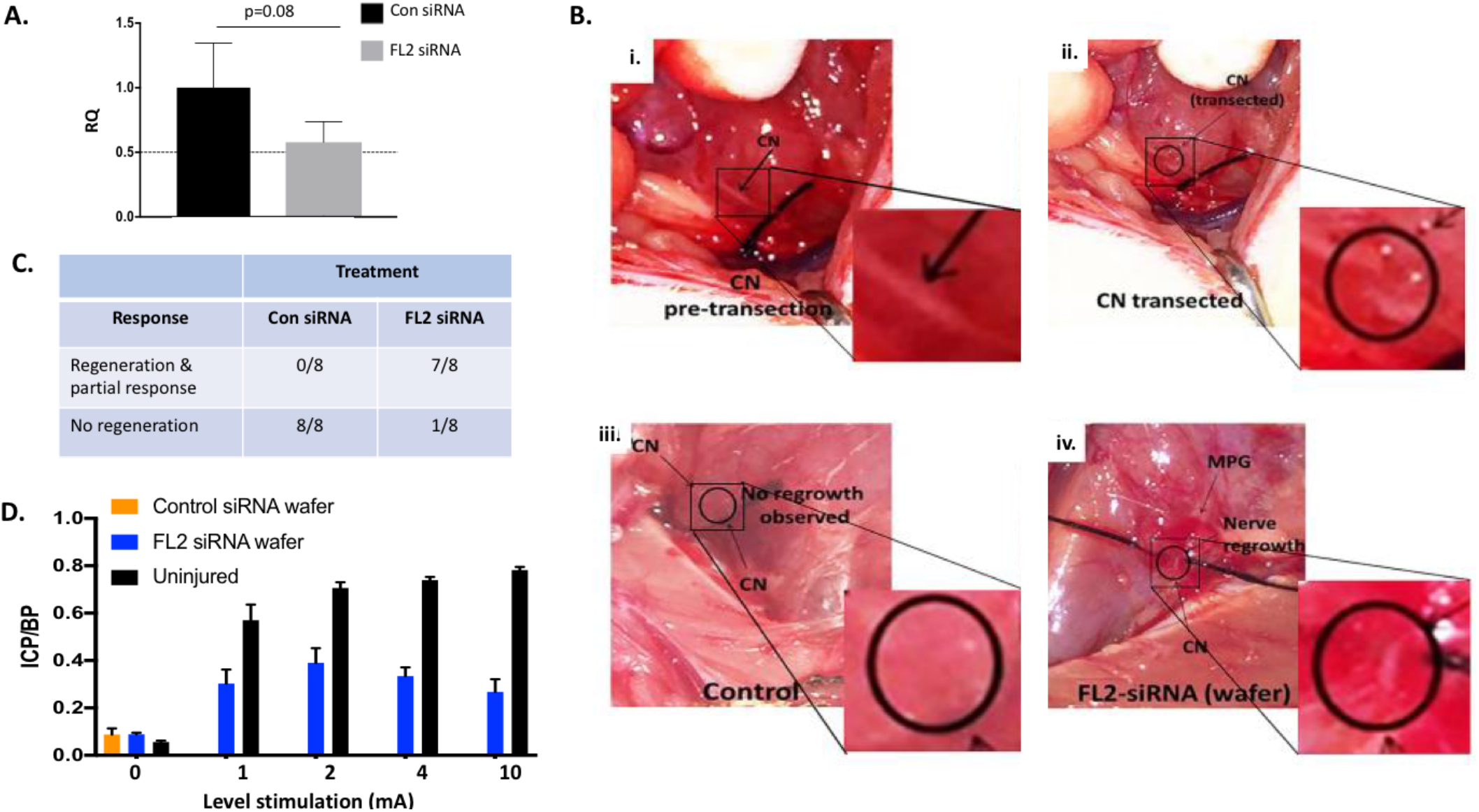
Application of FL2-siRNA after bilateral cavernous nerve transection leads to visible regeneration and partial recovery of erectile function. **A)** Relative levels of FL2 mRNA in the MPG two weeks after CN transection and wafer treatment (p = 0.08, n = 5). **B)** Images of the CN prior to transection (i); immediately after transection (ii); and 2 weeks post transection and Con siRNA or FL2 siRNA wafer treatment (iii and iv respectively). **C)** Table showing the number of animals which showed visible regeneration and partial recovery of erectile response and those that showed no regeneration; FL2 siRNA treatment promoted regeneration in 7 out of 8 animals whereas 0 control animals showed regeneration (Fishers exact test, P-value = 0.0004). **D)** Mean maximal intracorporal pressure/blood pressure (ICP/BP) measurements following different levels of stimulation of transected/ siRNA wafer treated animals and naïve age-matched controls (mean ± S.E.M.). Note control siRNA wafer transected nerves could not be stimulated due to the degree of retraction of the severed nerve segments, therefore only baseline ICP/BP is shown.

Two weeks following treatment of CN-transected animals with FL2-siRNA-wafer, 7 out of 8 animals exhibited visible CN regeneration, whereas in animals treated with control-siRNA-wafers, the two severed nerve segments had further retracted from each other and there was no visible nerve regrowth (Fishers exact test, p = 0.0004, Figs. 4B, C). In animals with visible regeneration of the CN, cavernosometry was performed (as depicted in supplementary Fig. S7B and C). Cavernosometry could not be performed on any control animals since there was no nerve to stimulate. At all levels of electrostimulation, animals treated with FL2-siRNA-wafer had a significant erectile response, as evidence by an increase in the intracorporal pressure relative to systemic blood pressure (ICP/BP) over basal (unstimulated) ICP/BP (Fig. 4D), indicating regenerated axons had reached the target tissue. The maximal ICP/BP reached was at the 2 mA level of stimulation, and was 69% of the erectile response achieved when naïve age matched control animals were stimulated at the same level. At higher levels of stimulation than 2 mA, both the maximal erectile response and percentage response relative to naïve animals decreased.

A segment of (unstimulated) regenerated nerve distal to the transection site was harvested from an FL2-siRNA wafer treated animal 4 weeks following transection and treatment, along with a distal nerve segment from an uninjured control animal. The nerves were processed for ultrastructural analysis by transmission electron microscopy (TEM) (Fig. 5). Large diameter myelinated axons as well as remak bundles of small diameter unmyelinated axons were identified in cross sections of regenerated nerve tissue (though, as would be expected, the remak bundles were sparser and contained fewer axons than those observed in the uninjured nerve at this timepoint). It was not possible to harvest a control-siRNA-treated nerve segment distal to the injury site because of the degree to which the nerve segment had retracted.

**Figure 5.**
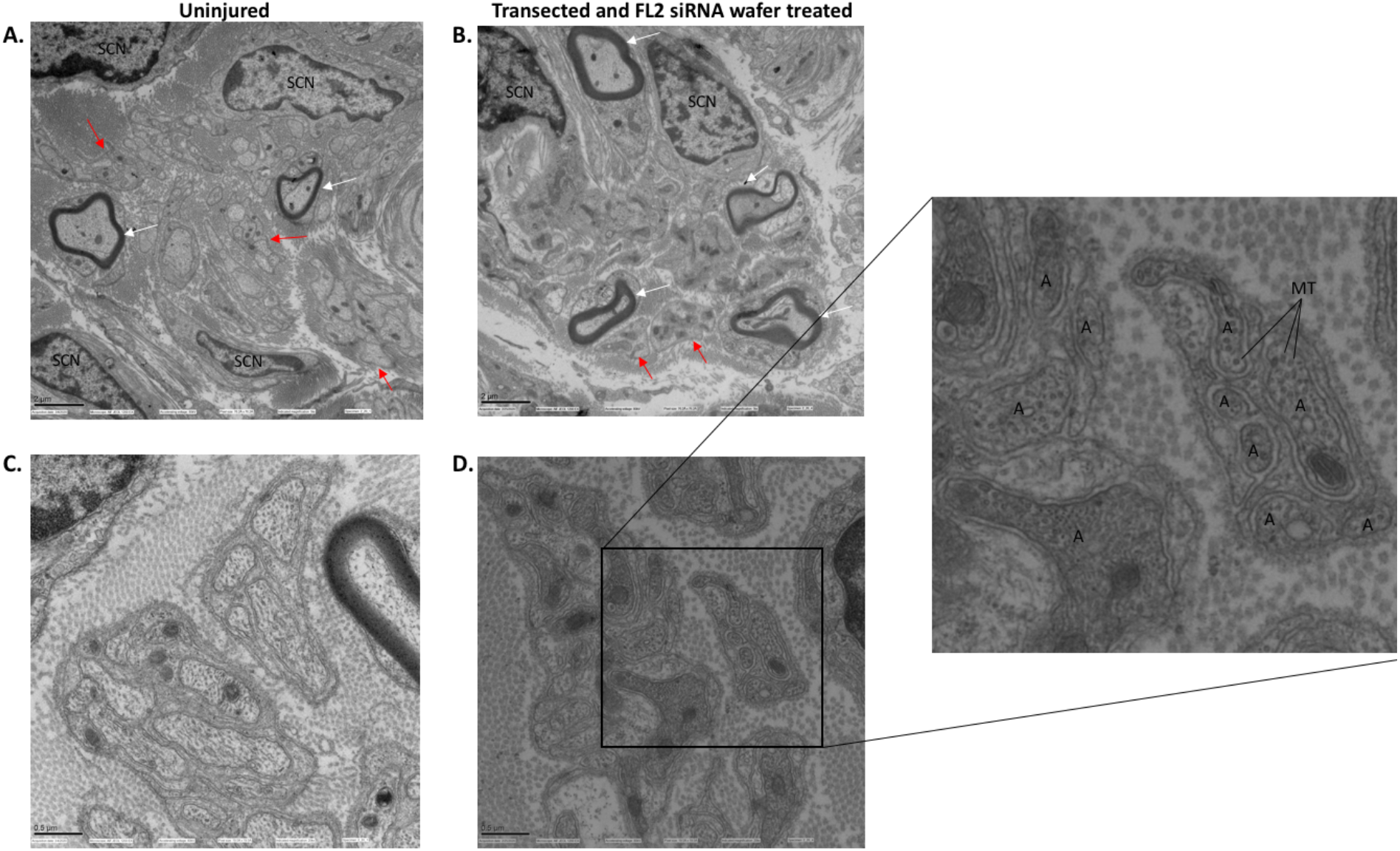
Regenerated myelinated and unmyelinated axons are present in the CN distal to the injury s ite following transection and FL2-siRNA wafer treatment. **A, B)** Low magnification transmission electron microscopy (TEM) images of a distal segment of CN from an uninjured animal and one from a transected and FL2-siRNA-wafer treated animal, respectively, 4 weeks after transection and treatment (harvest of control-siRNA-wafer treated distal nerve segments was not possible due to retraction of the distal nerve segment). White arrows point out myelinated large diameter axons. Red arrows point to some of the remak bundles of unmyelinated small diameter axons. Scale bar = 2 μm. **C, D)** Higher magnification TEM images of an uninjured (C) and transected and FL2 siRNA wafer treated nerve (D). In the inset of D, “A” labels individual axons. Some individual MTs (which appear as hollow rings in the axon cross sections) are pointed out. Scale bar = 0.5 μm.

### FL2 knockdown after CN crush increases the density of nitrergic neurons in the caudal MPG and smooth muscle cell density in the corpora

To confirm the regenerative effect of down-regulating FL2 and gain further insight into the recovery of erectile function mediated by knockdown of FL2 expression, we investigated the expression of markers known to be correlated with erectile function, such as neuronal nitric oxide synthase (nNOS) positive neurons in the MPG [45, 46] and expression of nNOS and smooth muscle actin (SMA) in corporal tissue [47, 48] using a nerve crush model. The nerve crush model is arguably a more clinically relevant CN injury model, since transection of the CN is usually avoided in RP, whereas damage to the nerves in the form of traction or crush is more likely to occur. In these experiments, siRNA was delivered to the MPG and injury sites via suspensions of nanoparticle-encapsulated FL2-siRNA (FL2-npsi) or control-siRNA (control-npsi), after CN crush (the nerve was crushed for 2 minutes using smooth forceps, 2-3 mm from the MPG). A time-course for functional CN regeneration was determined by cavernosometry performed weekly over the course of a month. At the 3- and 4-week post-injury time-points, cavernosometry demonstrated the FL2-npsi treated animals had significantly improved erectile function compared to animals treated with control-npsi (Fig. S7A, B).

MPG and corporal tissue were harvested from animals treated with FL2-npsi or control-siRNA 4-weeks after nerve crush and treatment. Consistent with the improvements in ICP/BP observed with FL2-npsi treatment, we found a higher density of nNOS positive somas in the region of the MPG proximal to the CN in rats treated with FL2-siRNA compared to controls (Fig. 6A,B). An increase in nNOS levels in penile shaft was observed by western blot analysis (Fig 6C,D). In addition, FL2-npsi treated rats had significantly higher levels of SMA relative to controls in the penile shaft (Fig. 6C,E).

**Figure 6.**
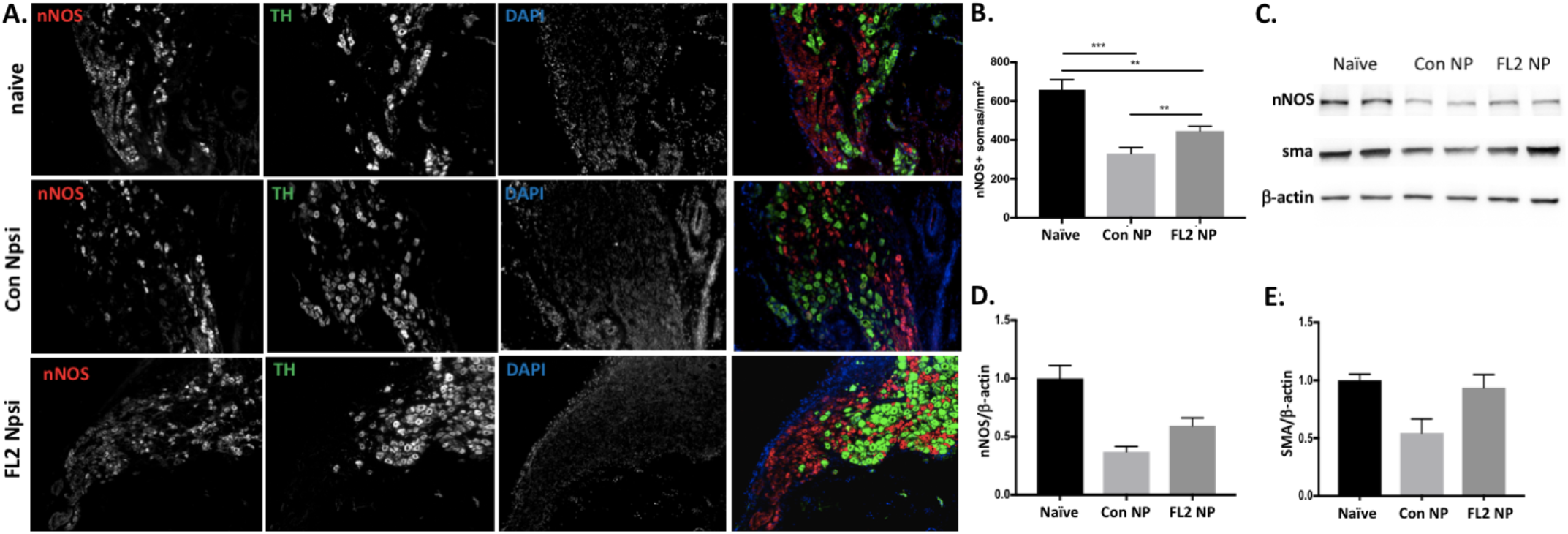
FL2 Npsi treated nerves had increased density of nNOS+ somas in the caudal MPG and increased nNOS and smooth muscle actin in the penis 4 weeks post cavernous nerve crush. **A)** Images of longitudinal sections of MPG proximal to the cavernous nerve from a naïve animal (top panel) and from animals 4 weeks after CN crush and treatment with control or FL2 NPsi (middle and bottom panels, respectively), stained for neuronal nitric oxide synthase (nNOS) to label nitrergic neurons, tyrosine hydroxylase (TH) to label sympathetic neurons, and DAPI. **B)** Average density of nNOS+ somas proximal to the CN in naïve, control and FL2 Npsi treated animals 4 weeks post CN crush. (Naïve: 655.2 ± 15.8; Con NPsi: 335.4 ± 34.7; FL2 Npsi: 451.7 ± 30.9. Mean ± SEM, **p<0.01, student’s t-test. n = 3 − 4 MPG per treatment group). **C)** Western blot of penile shaft samples from naïve animals and from rats 4 weeks after 4 minute CN crush and treatment with control of FL2 NP, probed for nNOS and smooth muscle actin (SMA). **D)** Relative levels of nNOS in the penile shaft by analyzed by western blot and normalized to β-actin, 4 weeks after 4 minute CN crush and treatment with FL2 or control NPsi (Con Npsi samples had a 64% reduction in nNOS levels compared to naïve animals, while FL2 Npsi samples had a 45% reduction, naïve n = 3, control Npsi n = 2, FL2 Npsi n = 3, blots run in duplicate). **E)** Relative levels of SMA in the penile shaft analyzed by western blot and normalized to β-actin, 4 weeks after 4 minute CN crush and treatment with FL2 or control NPsi (Con NPsi had a 45 ± 12% decrease relative to naïve animals, while FL2 Npsi sample SMA levels were comparable to naïve animals; naïve n = 3, control Npsi n = 2, FL2 Npsi n = 3, blots run in duplicate).

## Discussion

This is the first report identifying FL2 as a novel negative regulator of axonal growth and a mediator of growth cone steering in adult DRG neurons. Our data suggest that the enzyme suppresses axonal growth and mediates growth cone steering through its presumptive severing activity in the distal axon. Furthermore, we have shown *in vivo* that targeted knockdown of FL2 through the application of FL2-siRNA promotes peripheral nerve regeneration and recovery of nerve function.

The average ratio of dynamic to total MTs in the distalmost segment of DRG neurites significantly increased with FL2 depletion, indicated by the increase in tyrosinated to βIII tubulin and the corresponding decrease in acetylated to βIII tubulin. This observation is consistent with other studies which indicate that a more dynamic MT array in the distal axon is conducive to axon growth. MT deacetylation has been shown to be critical to the regeneration of adult peripheral neurons after injury (such a shift does not occur in injured CNS neurons, which lack the regenerative capacity of their PNS counterparts) [32], and dynamic MTs are critical for reformation of a viable growth cone [2, 49–51]. Notably, depletion of fidgetin—FL2’s closest structural relative—increases labile MT density in the axon and has also been shown to enhance axonal growth [19]. Further work is required to fully understand how FL2 remodels the axonal MT array and whether fidgetin and FL2 have overlapping roles within the axon or negatively regulate axonal growth through distinct mechanisms.

The fact that FL2 depletion attenuates the effects of inhibitory cues on growth cone advancement indicates the enzyme could be a promising therapeutic target for promoting axon growth in the CNS as well, where a major impediment to axon regeneration is the abundance of inhibitory molecules at the injury site [51, 52]. Axons in which FL2 has been depleted may be better able to traverse the glial scar after CNS injury. Less attention has been given to the role of inhibitory molecules in PNS regeneration, but inhibitory environmental cues, like sema3A, influence axon regeneration in the PNS as well [40, 41, 53]. In fact, Nangle and Keast demonstrated that adult parasympathetic and sympathetic neurons from the rat pelvic ganglia exhibit growth cone collapse in the presence of sema3A [40], raising the possibility that sema3A may play an important role in regeneration of pelvic ganglia neurons, but no studies testing this hypothesis *in vivo* have been conducted.

Remarkably, our nerve injury studies demonstrate that within two-weeks, targeted knockdown of FL2 following CN transection promotes functional nerve regeneration across a 3-4 mm gap between severed nerve ends, such that electrostimulation of the nerve elicits an erectile response. It was possible to observe visible regeneration of the CN, which, when examined at the ultrastructural level, had myelinated axons and Remak bundles distal to the site of injury. To our knowledge, regeneration across a gap of several millimeters following bilateral CN transection has not been observed without the use of a nerve graft. In studies where autologous and acellular nerve grafts were used to bridge 5 mm CN excision gaps in rats, the maximal ICP/BP achieved was 20-25% of that of naïve animals at one month, and approximately 50% at 3 months after surgery [54]. In our study, the average maximal ICP/BP reached in FL2-siRNA treated animals was 69% that of naïve age-matched control animals at 2 weeks, indicating a level of regeneration comparable and possibly superior to that seen with nerve grafts (albeit differences in the injury models and the time points at which nerve function was evaluated limit direct comparison).

The increase in nNOS positive somas in the MPG and corresponding increase in nNOS in the penile shaft provides further evidence of the regenerative effect of FL2 siRNA treatment. nNOS is present in the axons of nitrergic CN neurons innervating the penis, where it synthesizes nitric oxide (NO). NO release from the axons into the corpus cavernosum is the primary initiator of the erectile response [55]. The increased levels of SMA in the penile shaft observed with FL2 siRNA treatment are also indicative of improved CN function. CN injury and subsequent denervation of the corporal tissue causes apoptosis of smooth muscle cells and fibrosis in the corporal tissue [56, 57]. Loss of smooth muscle and increased fibrosis can permanently alter the fiberelasticity of the penis, impairing erectile response [58–60].

From a mechanistic standpoint, while our cell biology studies identify FL2 as a negative regulator of axonal growth, the regenerative effect of FL2 depletion on the injured CN is likely multifactorial (and not the consequence of FL2 downregulation in the axons alone). Indeed, the reformation of the neurolemma across the 3-4 mm gap between the transected nerve segments indicates a mechanism at work beyond accelerated axonal growth. Peripheral nerve repair is largely dependent on Schwann cells, fibroblasts, and immune cells, among others [61–63]. Following transection, axons distal to the injury site degenerate and the debris is cleared from the tissue by Schwann cells and immune cells in a process called Wallerian degeneration, a process which is critical for proper axon regeneration to ensue [61]. Meanwhile, Schwann cells must directionally migrate out of the nerve stumps and into the gap between the transected nerve ends, where they form corridors that guide regenerating axons into the distal nerve stump (referred to as Büngner bands) [64]. The migration of Schwann cells and the formation of these corridors across the nerve bridge is orchestrated by cell signaling between the Schwann cells and fibroblasts that accumulate in the gap following injury [62]. As FL2 knockdown was found to increase directional cell migration [20], it is possible that it improves formation of new nerve tissue across transection gaps by enhancing directional migration of Schwann cells and fibroblasts. Future work should determine the effects of FL2 downregulation in nonneuronal cells at the injury site, as well as further characterize molecular changes in the corporal tissue, CN, and MPG in order to elucidate the mechanism behind the observed regenerative effect.

## Methods

### Generation of the Fidgetin-like 2 conditional knockout mouse

The FL2 conditional knockout mouse was generated by GenOway. *Fignl2* is intronless, composed of 1 exon that extends over 4.2 kb on chromosome 15. Both ATG initiation and STOP codons are in exon 1. Polyadenylation signal consensus sequences can be found within the predicted 3’ UTR region (2273 bp). No cDNA or EST related to *Fignl2* gene is described in databases. However, the NCBI database proposes one sequence predicted by automated computational analysis for the *Fignl2* gene, in which a noncoding 5’UTR exon is present. The putative intron 1 would be 24kb large. Based on a bioinformatics analysis, a targeting strategy was designed to minimize the risk of interfering with regulatory elements while inserting exogenous elements and to avoid deregulating the targeted gene. An IRES-tdTomato sequence was inserted after the stop codon in exon 1 and two *lox*P sites flank a region starting upstream of the exon 1 coding sequence and finishing after the reporter (Figure S1A). After knock-in, a heterozygous conditional-knock-outline was generated. Breeding were established with C57BL/6 Flp deleter mice to excise the Neomycin selection cassette and to generate heterozygous mice carrying the neo-excised conditional knock-out allele. Figure S1B illustrates the Flp excision event. Heterozygous reporter floxed animals were produced, and the progeny genotyped by PCR (see Figure S2). The recombinase mediated excision event was then further validated by Southern blot on a subset of the PCR-positive animals (Figure S3). Through multiple rounds of breeding, a homozygous FL2 tdT flox/flox colony was established and confirmed by both traditional PCR based genotyping and automated qPCR based genotyping (Fig. S4 and S5).

### Preparation of PDL/laminin coated coverslips for neuronal culture

12 mm round #1 coverslips were cleaned with 10% HCl overnight, and then washed in an ultrasonicator in distilled and deionized water for 3 times (20 minutes per wash). The coverslips were then stored in 70% ethanol, and air-dried in the culture plate before coating procedure. 200 μl of 100 μM poly-D-lysine and 10 μM laminin was applied to the dried coverslips and the culture plate incubated at 37 C for 1-2 hours. Coverslips were then washed 3 times in sterile PBS.

### Neuronal cell culture

Dorsal root ganglia (DRGs) at the thoracic and lumbar levels were harvested from adult FL2-IRES-tdtomato flox mice between 6-12 weeks of age, and stored in media (MEM, 5% FBS, 1% glutamax, 500 units penicillin and 500 μg of streptomycin) on ice. DRGs were cleaned (extra tissue removed), washed in MEM, and then digested in 1 mg/ml of Collagenase A (Roche, 10103578001) in MEM for 90 minutes at 37 C, followed by TrypLE (Invitrogen) for 15 to 20 minutes at 37 C as previously described [65]. The DRGs were then washed with warm media and triturated in 1 ml of media with a P1000 pipette to prepare a single cell suspension in media. The cell suspension was transferred to the top of 2 ml of warmed 15% BSA in media, and spun at 160g for 10 minutes to clear myelin debris. The pellet was resuspended in fresh media with supplements (20 μM fluoro-deoxyuridine, 20 μM uridine, and 100 ng/ml nerve growth factor (NGF)). Cells were seeded onto PDL/LAM coated 12 mm coverslips in a 24 well plate at a concentration of roughly 10,000-15,000 neurons per coverslip. Neurons were treated with AV containing either a GFP encoding plasmid or a Cre recombinase encoding plasmid (Vector Biolabs, #1060 and #1045) at a viral titer of approximately 133-200 viral particles/cell (0.2 μl of a 1×10^10^ pfu/ml stock per well, in 300 μl of media). One week after transduction with the virus, neurons were replated onto either PDL/laminin coated coverslips or PDL/laminin coated aggrecan stripes for neurite growth to being anew. For the replating, neurons were dislodged by gentle pipetting 10 times with a P1000 pipette. This method dislodges neurons, while leaving nonneuronal cells attached to the coverslip, so that the replated suspension is very clean. The total suspension of dislodged cells from one well was replated onto a new coverslip. The concentration of resuspended cells prior to replating was too low to accurately quantify with a hemacytometer, but similar densities between replated GFP AV and Cre AV treated neurons was confirmed after replating and again after fixation and staining. For the rat FL2 knockdown experiments in Figure S6, the same dissection and culturing protocol was followed, except cells were transduced with AAV5 containing a control scrambled shRNA plasmid or a FL2 shRNA plasmid (Vector Biolabs). shRNA sequences were cloned into a D2-U6 vector. Scrambled sequence: CAACAAGATGAAGAGCACCAACTCGAGTTGGTGCTCTTCATCTTGTTGTTTTTG; FL2 sequence: CACCGCTGGAGCCCTTTGACAAGTTCTCGAGAACTTGTCAAAGGGCTCCAGCTTTT.

### Immunocytochemistry

Neurons were rinsed in pre-warmed HBSS and then fixed in warm 4% paraformaldehyde, 0.15% glutaraldehyde, 0.1% triton X 100, in BRB80 (80 mM PIPES, 1 mM EGTA, 1 mM MgCl_2_, pH 6.9) at 37 C for 10 minutes and then room temperature (RT) for 10 more minutes. Cells were rinsed twice PBS, incubated with 5-10 mg/ml sodium borohydrite in PBS for 15 minutes, rinsed and then permeabilized with 0.4% triton X 100 in PBS for 5 minutes. Cells were rinsed with PBS 3 times and incubated in blocking buffer (5% normal goat serum, 0.2% sodium azide, 0.1% triton X 100 in PBS) for 1 hour at RT or overnight at 4 C. Primary antibodies diluted in blocking buffer for applied for 2 hours at RT. After three 5 minute washes in PBS + 0.05% tween (PBST) secondary antibodies were applied for 1 hour at RT, then coverslips washed and mounted. Primary antibodies used: rabbit anti-βIII tubulin (Biolegend); anti-mouse tyrosinated tubulin (Abcam); anti-mouse acetylated tubulin (Abcam). Secondary antibodies used: Cy5-anti-rabbit (Jackson Immunoresearch), AF-568 anti-mouse (Thermo Fisher Scientific). Actin was stained with phalloidin-AF 488 (Thermo Fisher Scientific).

### Imaging and Analysis

Neurons were imaged using an EVOS Auto FL epifluorescent microscope (Life Technologies) at 10 X magnification for neurite length analysis and aggrecan stripe assay analysis, and 40X magnification for quantitative IF and growth cone morphology experiments. For neurite length analysis, the semi-automated ImageJ plug-in NeuronJ was used. The length of the longest neurite was measured to quantify average axon length. All neurite-bearing neurons where the longest neurite was clearly discernible and traceable were included in the analysis. To quantify the ratio of tyrosinated: βIII tubulin, the two fluorescence channels were combined into a merged RGB image in ImageJ and a line drawn over the distal neurite starting from the growth cone tip and moving toward the soma. RGB profiler was then used to obtain the intensity values of the two channels over each pixel along the line. These values were background corrected and then the ratio calculated along the line. To calculate the ratios of acetylated:βIII tubulin, ROIs were created around the distalmost 50 μM of neurites as well as the proximal-mid sections of neurites in ImageJ, and the total fluorescence in the separate channels was measured for each RO1, background corrected, and then the ratio calculated. Neurites from a minimum of 28 neurons were analyzed per treatment group per experiment. Experiments performed in triplicate.

### Aggrecan Stripe Assay

Stripes were prepared as previously described [37] with minor modifications. Briefly, aggrecan and alexa-fluor anti-mouse 568 were mixed in sterile PBS to final concentrations of 0.2 mg/ml aggrecan and 50 μg/ml AF-568 anti-mouse antibody. A Hamilton syringe was used to inject the mixture into zig-zag stripe silicon matrices (purchased from Prof. Martin Bastmeyer). The injected matrices were incubated at 37 C for 40 min., then 300 μl of PBS was washed through the matrices 3 times, after which the silicon matrices were removed from the dish. The stripe area was then incubated with PDL/laminin solution at 37 C for 60 min. The stripe areas were washed with PBS, and then culture media applied to the area until immediately before neurons were seeded onto stripes. The presence of CSPG on the stripes was confirmed by staining with an a-CSPG antibody, 3 days after incubation with media. For the experiments, AV-treated neurons were fixed and immunostained for βIII tubulin. Stripes and neurons were imaged at 10X on an EVOS Auto-FL microscope (Life Technologies). For quantification of neurite crossings the following criteria were used: Any time an axon’s trajectory came within several μm of stripe and then angled away from the stripe to stay within the growth permissive area, this was counted as a GC turning event. If it was unclear whether a neurite responded to a stripe or not (i.e. if the neurite grazed the side of a stripe boundary crossing into it briefly but continuing on its trajectory back into the permissive stripe zone), this neurite was not included in the total neurite count.

### Sema3A-induced growth cone collapse analysis

Neurons were replated as describe above. Two days after replating, media was changed to media with 100 ng/ml recombinant mouse Sema3A-Fc (R&D) or Fc (R&D) as a control for 2 hours, then fixed and immunostained for βIII tubulin and actin. Growth cones (GCs) were imaged at 40X magnification. For growth cones to be count as collapsed, growth cone MTs had to be bundled and no filopodia present (the width of these collapsed GCs was less than 2 μm). Over 100 neurites from 28-32 neurons were analyzed per condition each experiment. Experiment was performed in quadruplicate for AV treated neurons and in triplicate for non-AV treated control neurons. All imaging and analysis for this experiment was performed blind.

### siRNA nanoparticle synthesis

Nanoparticles were prepared as previously described [20] with minor modifications: 500 μl of tetramethyl orthosilicate (TMOS) was hydrolyzed in the presence of 100 μl of 1 mM HCl by sonication on ice for about 15 min, until a single phase formed. The hydrolyzed TMOS (100 μl) was added to 900 μl of 10 μM of pooled siRNA against rat FL2 (siRNA from Sigma-Aldrich: SASI_Rn02 00314854; SASI_Rn02 00314854; SASI_Rn02_ 00389576) or the negative control siRNA (Sigma, Universal Negative control B) solution containing 10 mM phosphate, pH 7.4. A gel was formed within 10 minutes. The gel was frozen at −80°C for 15 minutes and lyophilized. The dried sample was ground into a fine powder with a mortar and pestle. The nanoparticles were then resuspended in sterile PBS at an siRNA concentration of 10 μM, and stored at −80 C until immediately before use.

### Cavernous nerve injuries and treatments

Animals were anesthetized with ketamine and xylazine. A midline abdominal incision was made and the CN exposed and isolated. Three rat models of CN injury were used: mild (a smooth clamp applied for 2 minutes to the CN), moderate (a serrated clamp applied for 4 minutes to the CN) and severe (CN transection). In the case of the crush injury studies, 10 μl of a 10 μM control or FL2 Npsi suspension was applied directly onto the nerve injury site. For the transection model, a collagen chondroitin-sulfate hardened microgel (wafer) containing control or FL2 siRNA (synthesized by and purchased from BioAdd Laboratory at Stanford University) was applied directly to the wound. The approximate concentration of siRNA in the wafer was 13.3 μg siRNA/100mg of wafer, and ~15 mg of wafer was applied. The wafer completely covered the nerve injury site as well as the major pelvic ganglion.

### Cavernosometry

Animals were anesthetized with a Nembutal^®^ Sodium solution, 50mg/ml, and erectile function was determined by electrostimulation of the CNs and by measuring intracavernous pressure (ICP): through a repeat midline abdominal incision, the CNs were exposed and isolated. Cannulas were inserted into the crura at the base of the penis and the carotid artery to measure the ICP and BP, and a bipolar stainless steel electrode, inserted above the site of nerve transection, was used to directly stimulate the CN (probes 2 mm in diameter, separated by 1 mm) via a signal generator and a custom-built constant-current amplifier generating monophasic rectangular pulses with stimulus parameters ranging from 1.5-6 mA, 20 Hz, pulse width of 0.2 ms, and duration of 50 s. The ICPs and BP were recorded in all rats using a bioinformation acquisition system. The maximal ICP/BP ratios in the experimental animals were calculated for each level of stimulation.

### Quantification of FL2 mRNA expression by RT-qPCR

To analyze FL2 mRNA expression *in vitro,* RNA was isolated from cell cultures using Trizol (Fisher, X) following manufacturer’s protocol. 200-300 ng of RNA were reverse transcribed using the SSVilo IV kit (Invitrogen). To verify downregulation of FL2 mRNA following wafer treatment *in vivo,* rat MPG were harvested and kept in RNAlater (Qiagen) immediately after cavernosometry. MPG were homogenized using a bullet blender (Next Advance) and RNA reverse transcribed using High-Capacity cDNA Reverse Transcription Kit (Ambion) (500ng/reaction). PowerSybr green Master Mix was used for qPCR, using the 7300 Real-Time PCR system (Applied Biosystems). The following primer sequences were used: mouse Fignl2: GTTCACACTCCTCACACCTG and GCTCCTAGATCCCTTCATGTTC; rat Fignl2: GAGTTGCTGCAGTGTGAATG and CTCTGTGCTTCTGTCTCTGT; βactin (mouse and rat): CGTTGACATCCGTAAAGACC and TCTCCTTCTGCATCCTGTCA; tdtomato: CACGCTGATTCTACAAGGTGAA and CCCATGGTCTTCTTCTGCATTA. RPL19 was used as reference gene for the *in vivo* experiments (ACCCCAATGAAACCAACGAA and TCAGGCCATCTTTGATCAGCTT). Results were analyzed using the comparative 2^−ΔΔCt^ method. For adult DRG cultures, neurons from 4 mice were pulled into one RNA extract for each treatment groups.

### Transmission Electron Microscope Analysis

Distal nerve segments from one (unstimulated) regenerated FL2 siRNA wafer treated nerve and one uninjured control animal were harvested while the animal was anesthetized with ketamine/xylazine. The samples were fixed with 2% paraformaldehyde and 2.5% glutaraldehyde in 0.1 M sodium cacodylate buffer, postfixed with 1% osmium tetroxide followed by 2% uranyl acetate, dehydrated through a graded series of ethanol and embedded in LX112 resin (LADD Research Industries, Burlington VT). Ultrathin (80 nm) sections were cut on a Leica EM Ultracut UC7, stained with uranyl acetate followed by lead citrate and viewed on a JEOL 1200EX transmission electron microscope at 80kv.

### Immunohistochemistry

MPG were harvested after sacrifice (by CO2) and fixed for 20 hr in 4% pfa, transferred to 30% sucrose for 48 hrs, and then cryosectioned in 10 μm slices longitudinally. Sections were collected sequentially onto slides. 3 sections per MPG, 100-150 μm apart from each other, were used for quantification of nNOS+ somas. Sections were immunostained for nNOS (Santa Cruz SC-5302) and tyrosine hydroxylase (Abcam ab112) (as a counterstain to visualize non nNOS+ somas) as well as DAPI according to standard a IHC protocol. Slides were imaged on an EVOS Auto FL epifluorescent microscope (Life Technologies) using a 10X objective. nNOS+ somas in caudal region of the MPG proximal to the CN were quantified (areas ranged 0.25-0.5 mm^2^). If there was a tear in the tissue in the analyzed region, this area was subtracted from the total area. 3-4 MPG were analyzed. The average density of the 3 sections was calculated to provide a single value for each MPG, and this value was used in statistical analysis. Slide selecting, imaging and analysis for these experiments was performed blind.

### Corporal tissue homogenate preparation and western blot analysis

A 0.5 cm segment of the mid penile shaft from each rat were flash frozen after sacrifice. The samples were then minced and homogenized in lysis buffer (50 mM Tris, pH 7.5, 2 mM EDTA, 2 mM EGTA, 150 mM NaCl, 1% triton-X, and 10% vol./vol. glycerol) using a bullet blender (Next Advance), then protein concentration quantified by BCA assay. 30-50 μg of protein were loaded into 4-20% gradient gels for SDS-PAGE at 35 mA using the BioRad mini-protean system, then protein transferred onto a nitrocellulose membrane for 1 ½ hr at room temp. at 100 V. Blots were blocked in 5% milk in TBS, and then incubated overnight with primary antibodies against nNOS+ (Santa Cruz), smooth muscle actin (Santa Cruz), or β-actin, (abcam) and incubated with secondary antibodies conjugated to horse radish peroxidase for 1 hr at room temp. In the case of the smooth muscle actin blots, blots were stripped after probing for a-actin in mild stripping buffer (67 mM glycine, 0.1 SDS%, 1% tween 20, pH 2.2), and then probed for smooth muscle actin overnight. 3 naïve, 2 control Npsi, and 3 FL2 Npsi samples were run in duplicate blots. Blots were visualized using the iBright system and densitometry was performed in either ImageJ or using iBright software automated analysis.

### Study Approval

Animal experiments were performed according to the guidelines published by the Institute of Laboratory Animal Resources of National Research Council and animal care for this study was approved by the Institutional Animal Care and Use Committee of the Albert Einstein College of Medicine.

## Acknowledgements

We thank Xheni Nishku, Frank Macaluso, and, in particular, Leslie Gunther-Cummins of the Einstein Analytical Imaging Facility for their assistance in preparing and imaging nerve samples by transmission electron microscopy. We thank the Dr. Bin Zhou and his laboratory for the use of their cryostat. Funding for these studies was provided by: NIH RO1 GM109909; RO1 DK109314; T32 5T32GM007491; NYS Spinal Cord Injury Research Board contract C030166; and Sexual Medicine Society of North America 2016 and 2018 graduate student research fellowships.

## Author Contributions

L.B., K.D., and D.S. wrote the manuscript; L.B. designed and performed cell biology experiments, histology, western blot experiments, and analyses; M.T. performed all surgeries, cavernosometry, and ICP/BP analysis; G.V. performed qPCR of MPG tissue samples; R.C. contributed intellectually and executed early pilot studies using siRNA in rat neurons; A.K. contributed intellectually and bred the mouse; O.V. assisted with morphometric analysis of neurons; P.N. and J.F. synthesized nanoparticles; K.D. and D.S. conceived of the project.

## Supplemental Materials

**Figure S1:**
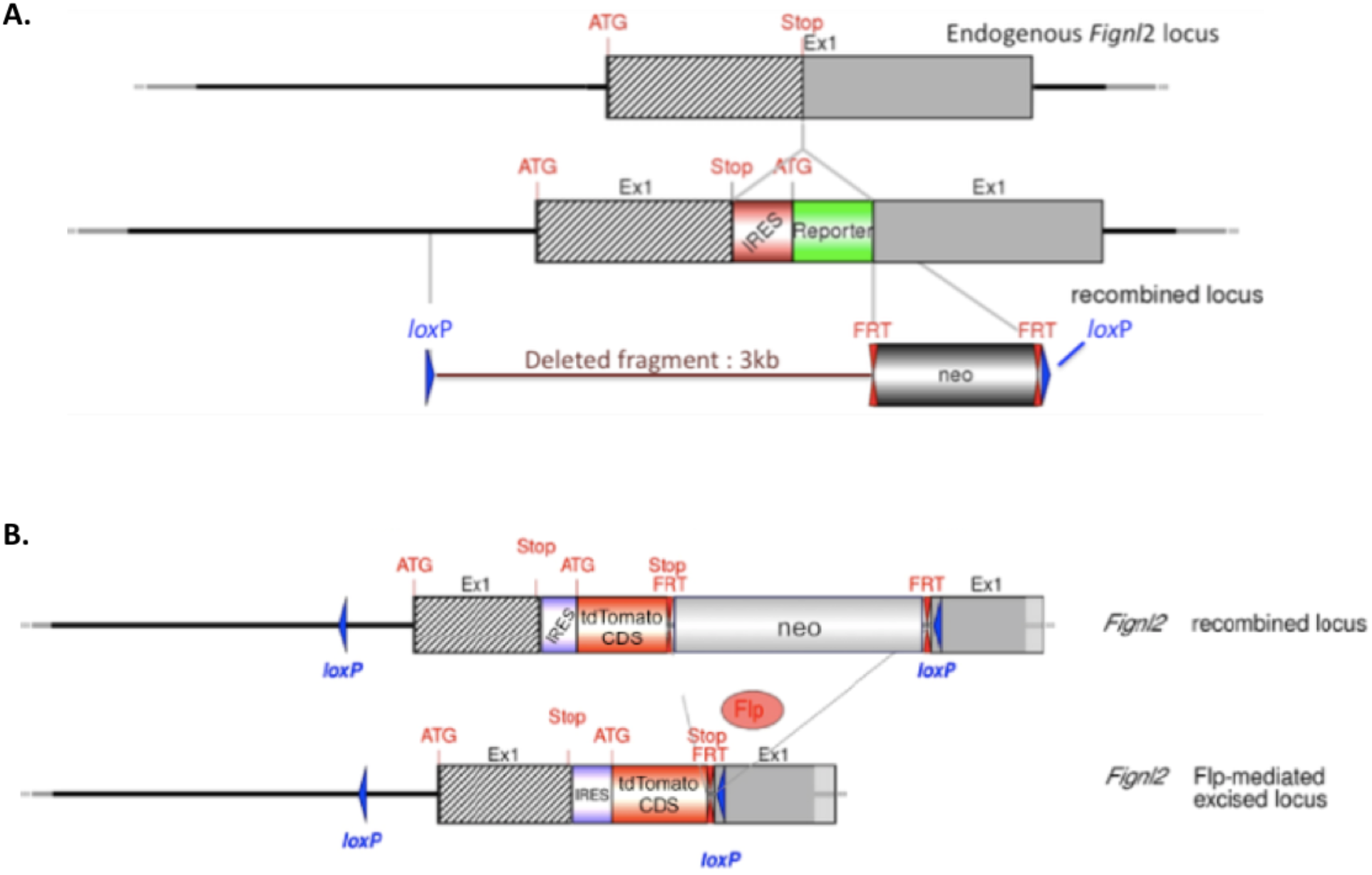
Insertion of loxP sites and IRES-tdTomato gene at FL2 locus. **A)** Schematic representation of the selected targeting strategy. Hatched rectangles represent Fignl-2 coding sequences, grey rectangles indicate non-coding exon portions and solid lines represent chromosome sequences. The neomycine positive selection cassette is indicated. loxP sites are represented by blue triangles and FRT sites by double red triangles. The initiation (ATG) and Stop (Stop) codons are indicated. The size of the flanked Fignl-2 sequence to be deleted is specified (3kb). **B)** Scheme of Flp-excision at the *Fignl2* recombined locus. Diagrams not depicted to scale.

**Figure S2:**
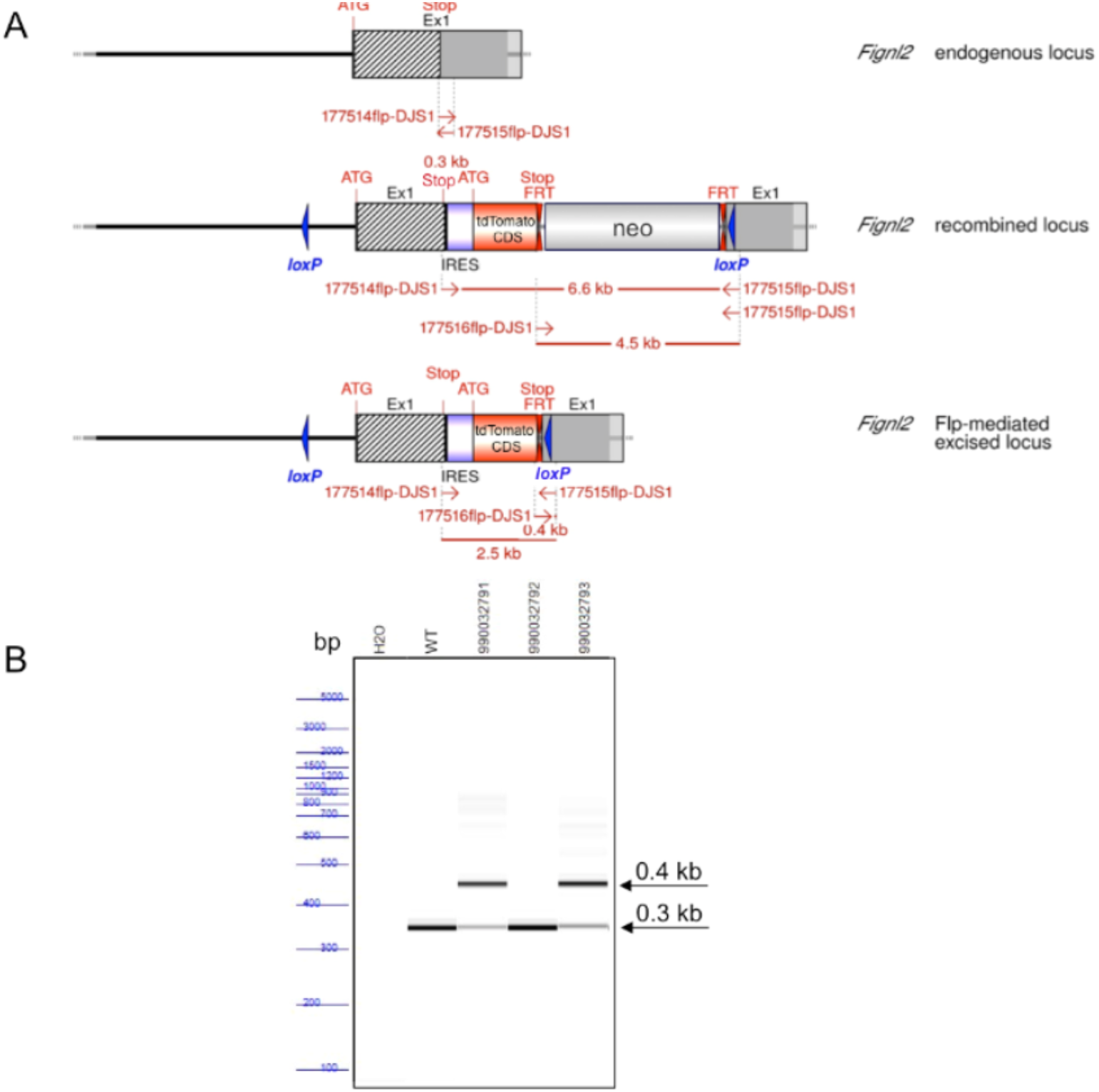
Flp-mediated excision of neomycin cassette. (A) Schematic representation of the Fignl2 wildtype, non-excised recombined and conditional Knockout alleles with the binding sites of the screening primers. (B) PCR using C57BL/6 wild-type genomic DNA (WT) was used as positive control. PCR without DNA as template (H2O) served as a negative control. PCR fragments were separated by capillary electrophoresis using AATI ZAGTM Fragment Analyzer and were analyzed using PROSize 2.0 analytical software. Note: Due to the large size of the 2.5 kb Flp-mediated excised allele PCR product, its amplification did not occur.

**Figure S3:**
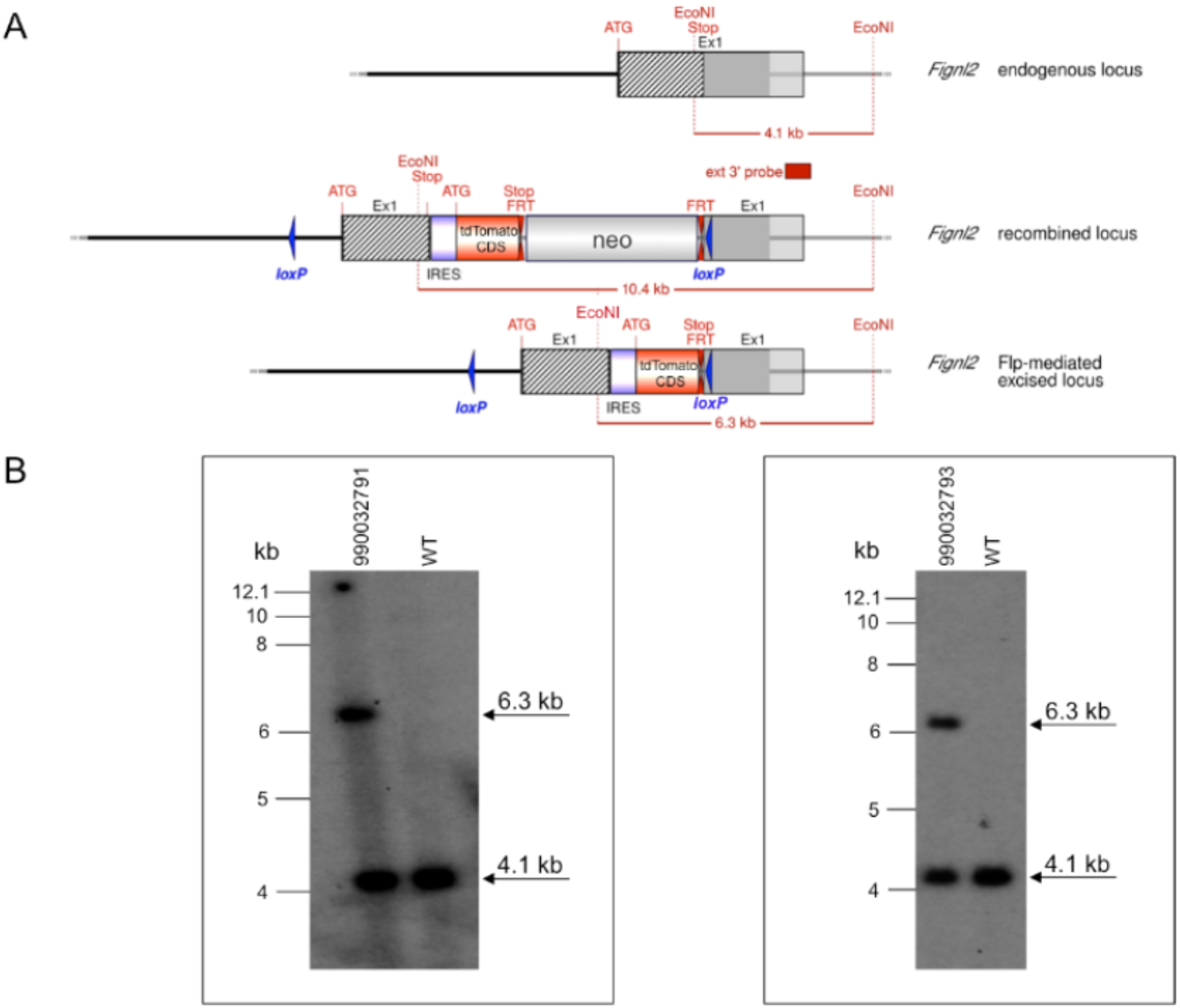
Representative Southern blot results of the animals of interest. (A) Schematic representation of the *Fignl2* wild-type, non-excised recombined and conditional Knockout alleles. (B) The genomic DNA of the tested animals was compared to C57BL/6 wild-type genomic DNA (WT).

**Figure S4:**
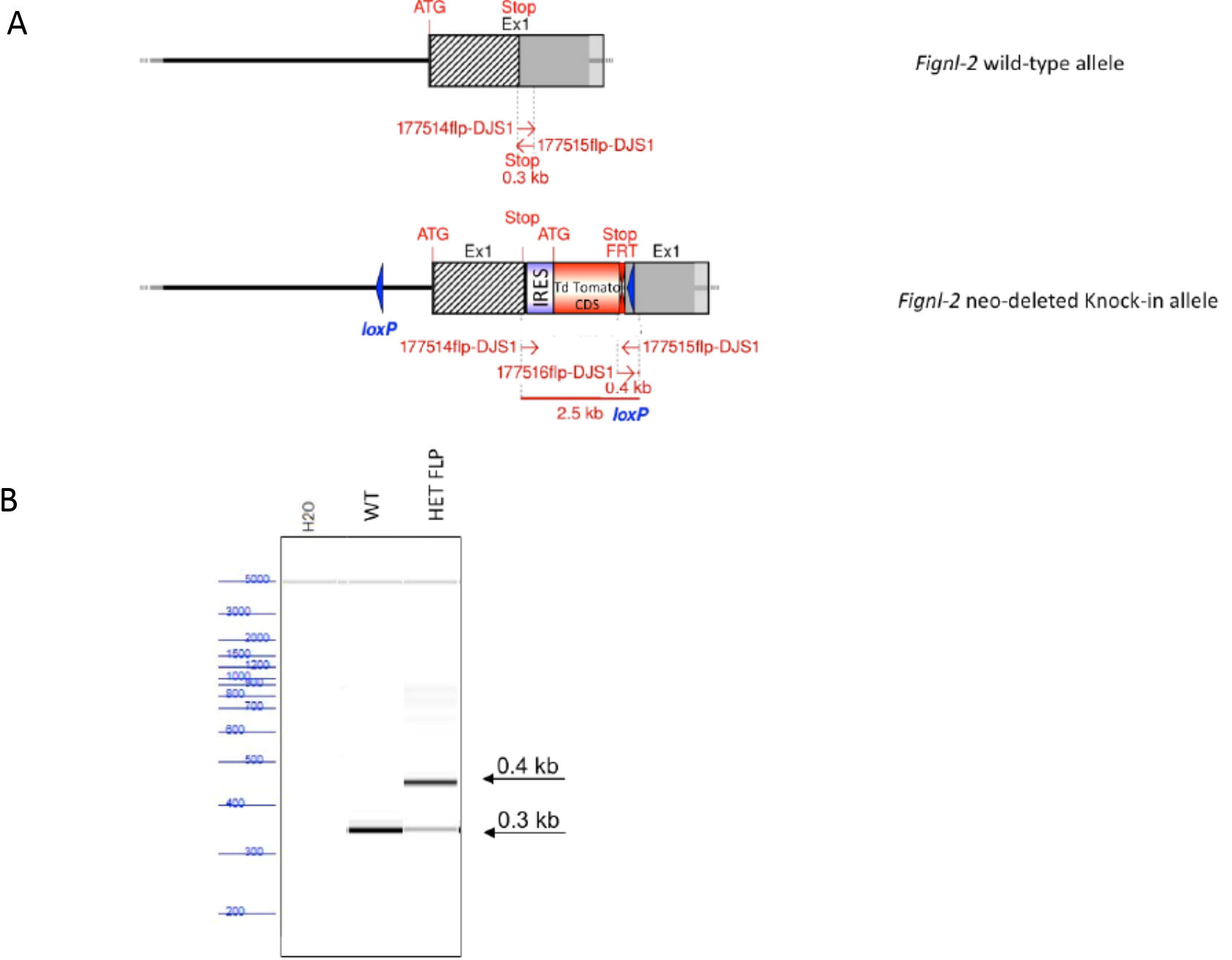
PCR identification of the FL2 neo-deleted knock-in allele. A) Schematic representation of the Fignl-2 wild-type and neo-deleted Knock-in alleles with the binding sites of the screening primers. B) The optimized PCR screening was conducted using genomic DNA of heterozygous conditional/ knock-out animals (HET FLP). PCR with C57BL/6 wild-type genomic DNA (WT) and without DNA (H2O) were used as positive and negative controls, respectively. PCR picture was obtained after loading the PCR reactions on the LabChip^®^ system from Caliper LifeSciences.

**Figure S5:**
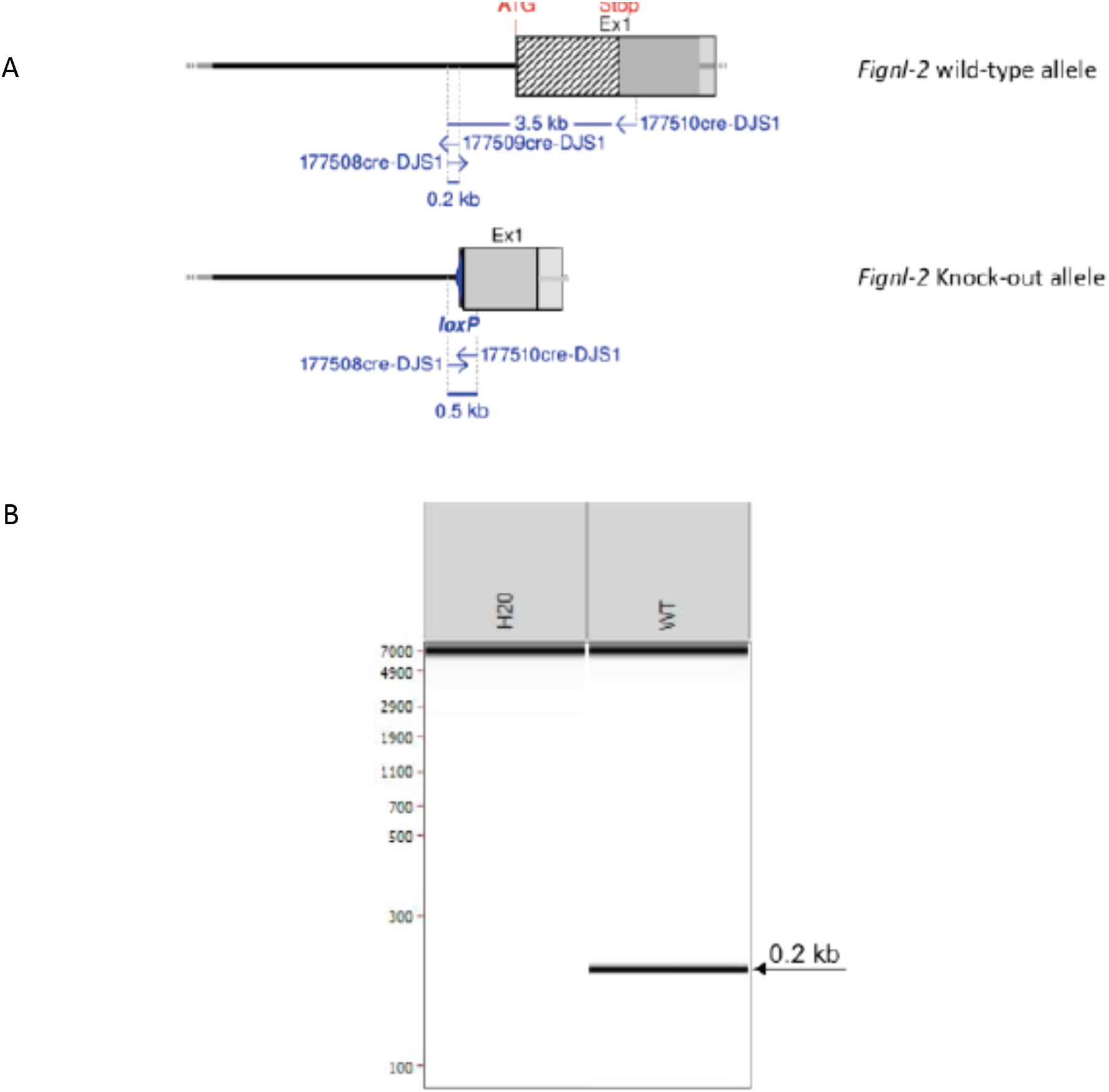
PCR identification of the Fignl-2 Knock-out allele. A) Schematic representation of the *Fignl-2* wild-type and Knock-out alleles with the binding sites of the screening primers. B) The optimised PCR screening was conducted using C57BL/6 wild-type genomic. PCR without DNA (H2O) was used as negative control. PCR picture was obtained after loading the PCR reactions on the LabChip^®^ system from Caliper LifeSciences.

**Figure S6.**
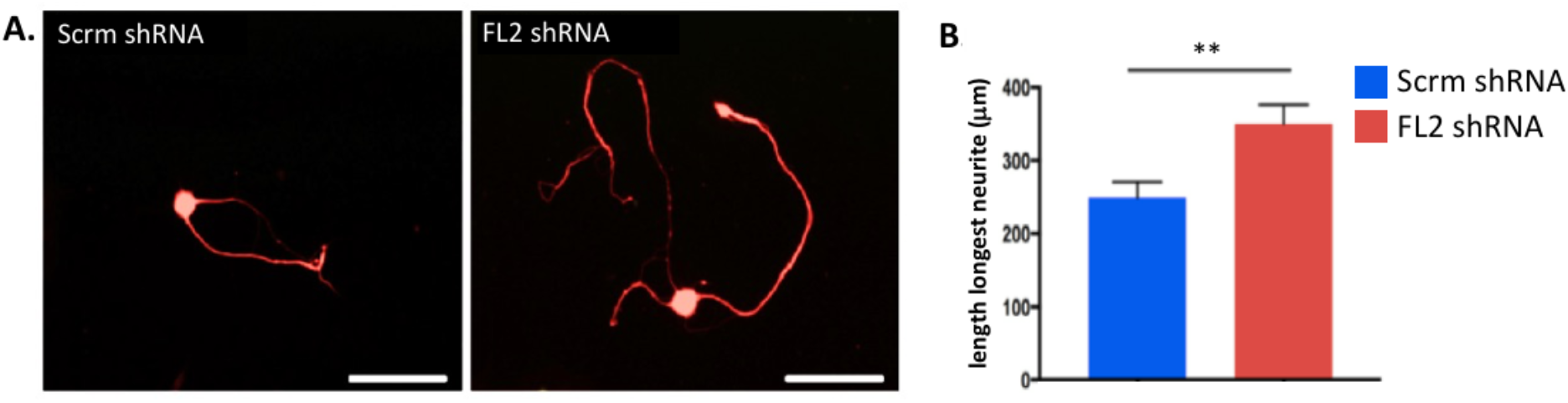
Knockdown of FL2 in rat DRGs promotes axonal regeneration. Preliminary study with adult rat dissociated DRG neurons, replated 5 days after transduction AAV5 containing with control (scrambled) shRNA or FL2 shRNA plasmids, and fixed 72 hours post replating. FL2 shRNA expressing neurons had significantly longer axons than control treated neurons. **A)** Immunomicrographs of replated neurons expressing Scrm shRNA or FL2 shRNA, 72 hours after replating. **B)** Graph of the mean length of longest neurites of Scrm shRNA and FL2 shRNA expressing cells (control shRNA n = 59, FL2 shRNA n = 68, student’s t-test, p < 0.01).

**Figure S7.**
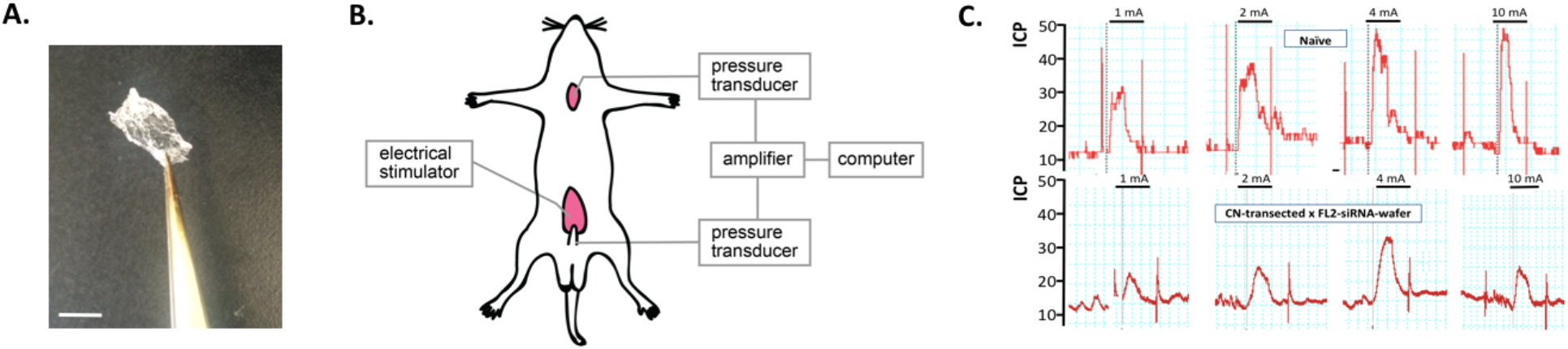
Assessment of erectile function after transection and treatment with siRNA wafers. **A)** Image of wafer embedded with siRNA prior to application onto wound. Scale bar = 0.5 cm. **B)** Schematic of cavernosometry set-up. Cannulas are inserted into the penile cruz and the carotid artery to measure the intracorporal pressure (ICP) and blood pressure (BP), respectively. A bipolar stainless steel electrode, inserted above the site of nerve injury, is used to directly stimulate the CN. **C)** Representative traces of the ICP in response to increasing levels of cavernous nerve stimulation in an uninjured animal and an animal that underwent bilateral CN transection and FL2 siRNA wafer treatment.

**Figure S8.**
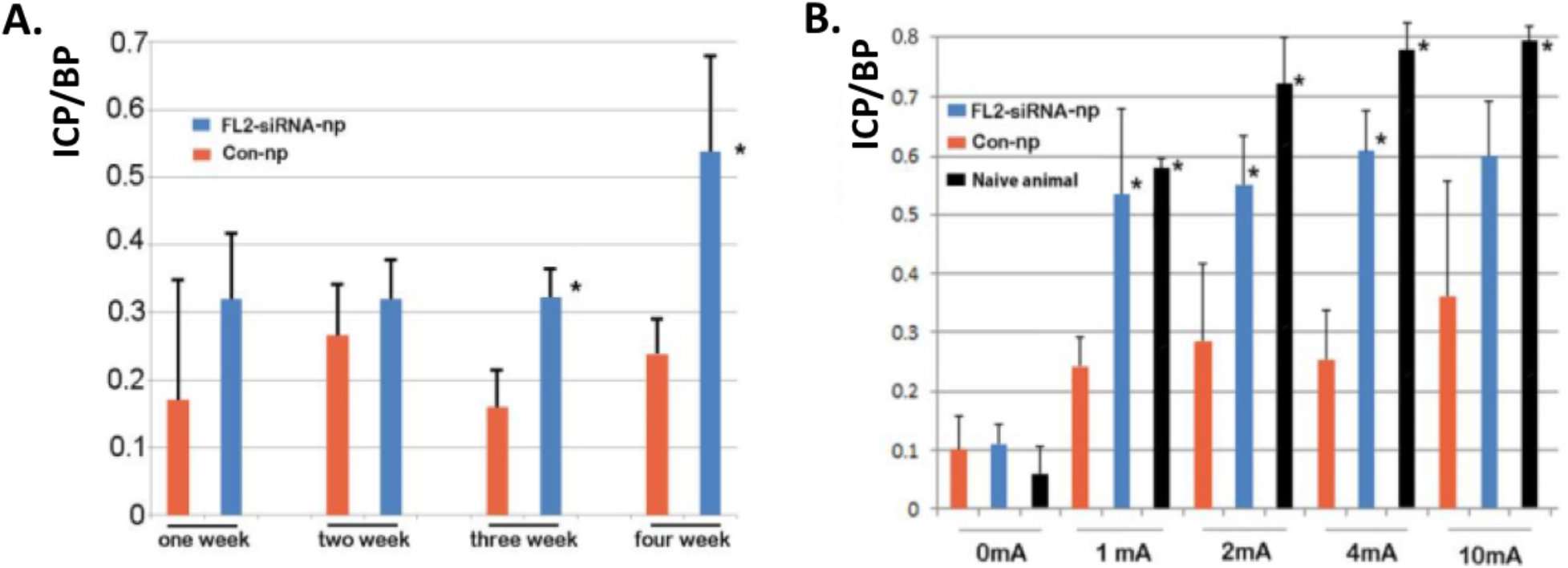
Treatment with nanoparticle-encapsulated FL2 siRNA improved erectile response at 3 and 4 weeks post nerve crush. **A)** Time course showing mean ICP/BP of control Npsi and FL2 Npsi treated nerves with 1 mA electrostimulation of the nerve (mean ± SD, *p<0.05, 5-6 animals per time point and treatment group). **B)** ICP/BP for naïve and NPsi treated rats 4 weeks after crush and treatment (mean ± SD, *p<0.05, n=6 per treatment group).

